# Neurobehavioural characterisation and stratification of reinforcement-related behaviour

**DOI:** 10.1101/778308

**Authors:** Tianye Jia, Alex Ing, Erin Burke Quinlan, Nicole Tay, Qiang Luo, Biondo Francesca, Tobias Banaschewski, Gareth J. Barker, Arun L.W. Bokde, Uli Bromberg, Christian Büchel, Sylvane Desrivières, Jianfeng Feng, Herta Flor, Antoine Grigis, Hugh Garavan, Penny Gowland, Andreas Heinz, Bernd Ittermann, Jean-Luc Martinot, Marie-Laure Paillère Martinot, Frauke Nees, Dimitri Papadopoulos Orfanos, Tomáš Paus, Luise Poustka, Juliane H. Fröhner, Michael N. Smolka, Henrik Walter, Robert Whelan, Gunter Schumann, IMAGEN Consortium

## Abstract

Reinforcement-related cognitive processes, such as reward processing, impulsiveness and emotional processing are critical components of externalising and internalising behaviours. It is unclear to what extent each of these processes contributes to individual behavioural symptoms, how their neural substrates give rise to distinct behavioural outcomes, and if neural profiles across different reinforcement-related processes might differentiate individual behaviours. We created a statistical framework that enabled us to directly compare functional brain activation during reward anticipation, motor inhibition and viewing of emotional faces in the European IMAGEN cohort of 2000 14-year-old adolescents. We observe significant correlations and modulation of reward anticipation and motor inhibition networks in hyperactivity, impulsivity, inattention and conduct symptoms, and describe neural signatures across neuroimaging tasks that differentiate these behaviours. We thus characterise shared and distinct functional brain activation patterns that underlie different externalising symptoms and identify neural stratification markers, while accounting for clinically observed co-morbidity.

## INTRODUCTION

Reinforcement-related behaviours are commonly implicated in normal behaviour and psychopathology. Symptoms of dysfunctional reinforcement-related behaviours may present as hyperactivity, inattention, conduct and emotional problems^1^, and substance use^2^. These symptoms are manifest in common psychiatric disorders, such as depression, ADHD, addictions, conduct disorder and psychosis ^3, 4^. Reinforcement-related behaviours involve similar cognitive processes, including reward processing, impulsiveness and emotional processing^5^. However, while such similar cognitive processes are involved in different disorders, there are clear differences in their behavioural presentation in each disorder. It is unclear if and how the cognitive processes involved in reinforcement-related behaviours are modulated to achieve the observed behavioural differences. Identifying the brain activity patterns related to various manifestations of dysfunctional reinforcement-related behaviour might aid in the characterisation of underlying biological mechanisms, and the identification of targets for therapeutic intervention^6^. Furthermore, clinically relevant psychiatric symptoms typically are characterised by dysfunctions not only in one but often in several reinforcement-related domains. For example, ADHD symptoms are known to involve dysfunctional executive control^1^, as well as dysfunctional reward processing^7^. We were interested in dissecting the contribution of different domains of reinforcement-related behaviour to distinct disorder symptoms, and thus characterise a profile of brain activation specific for each disorder.

Whereas animal models have identified networks of multiple cortical and subcortical brain regions involved in reinforcement-related behaviours^8^, analyses in humans are often based on a few pre-defined regions of interest (ROI). These include the ventral striatum (VS) and orbital frontal cortex (OFC) for reward processing^9^, right inferior frontal cortex (rIFC) for response inhibition^10^, and amygdala and superior temporal sulcus (STS) for emotional processing ^11, 12^. Often, the underlying assumption is that a cognitive process can be represented by a few key brain regions. However, we^13^ and others ^14-17^ have shown that task-induced brain activity may involve a complex network of cortical and subcortical brain regions. We do not know, however, how these networks relate to observable behaviour.

In this paper we provide a systematic characterisation of brain activity in reinforcement– related behaviour, measuring BOLD-response during tasks targeting reward anticipation, motor inhibition and socio-emotional processing. We compare their common and distinct brain activity patterns and assess the modulation of task-specific networks in externalising (e.g. hyperactivity, inattention, impulsivity and conduct symptoms) and internalising (e.g. emotional and anxiety symptoms) behavioural symptoms^18^. We also identify signatures of brain activity across tasks that best characterise symptoms of externalising disorders.

## RESULTS

### Summary of Analysis Strategy

We aim to compare brain activity during functional neuroimaging tasks measuring reward anticipation, motor inhibition and emotional processing of 1506 14-year-old adolescences from IMAGEN project^5^. We reduced the dimensionality of brain activation by applying a weighted voxel co-activation network analysis (WVCNA) ^13, 19^, followed by a hierarchical clustering analysis. The combination of both methods could efficiently reduce dimensionality while still preserving localised network features from WVCNA. We then calculated the overall correlation between fMRI clusters and symptoms of externalising or internalising behaviours using ridge-regularised canonical correlation analysis (RCCA)^20^, a method to detect multivariate relations between different data types.

First, we tested for an overall significant correlation of externalising or internalising symptoms with brain network activation across all fMRI tasks. In cases where we established an overall correlation, we looked for association of each fMRI network with externalising or internalising behaviours. Finally, we investigated the sensitivity and specificity of fMRI clusters across different behaviour components. The above workflow was illustrated as Figure 1.

**Figure 1.**
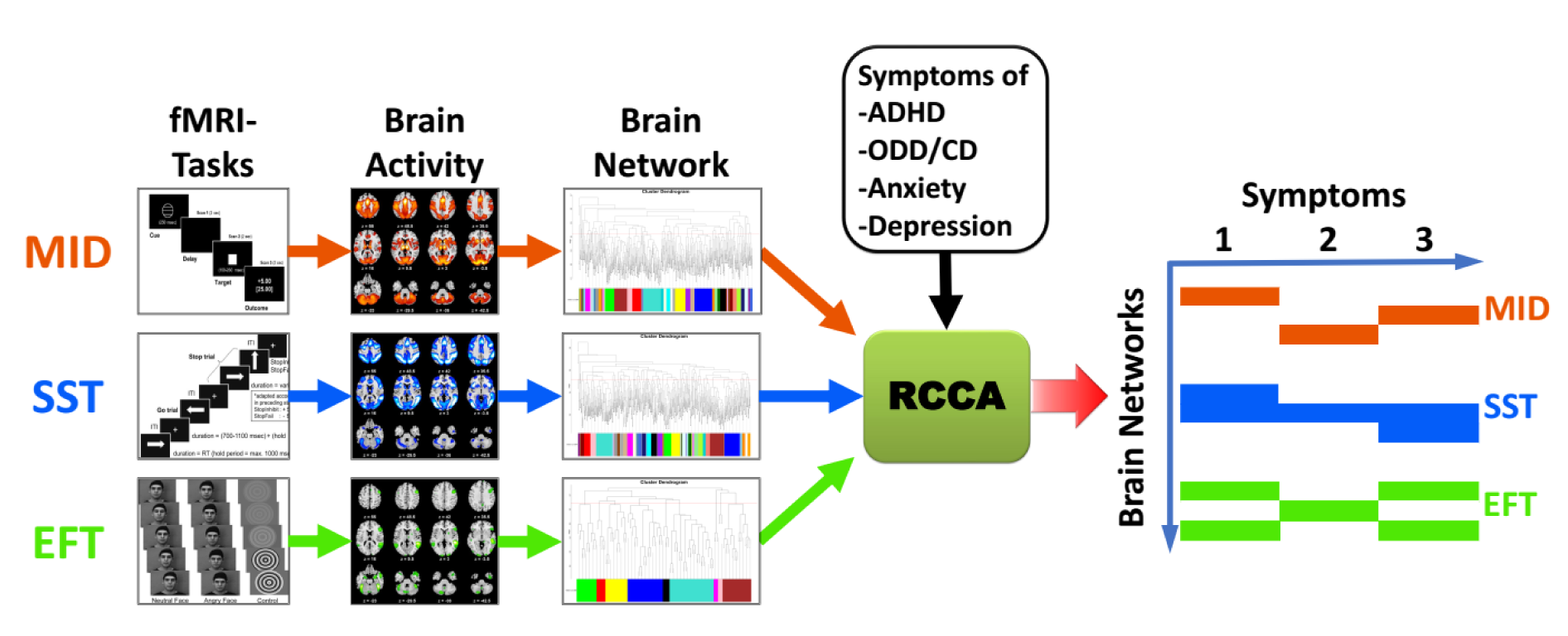
Workflow of the Analyses. We included the monetary incentive delay task (MID) as a measure of reward processing, the stop signal task (SST) as a measure of impulsivity (motor inhibition), and the emotional face task (EFT) as a measure of emotional processing. Only strong brain activation (Cohen’s D>0.30) was included in the analyses. The weighted voxel co-activation network analysis (WVCNA) in combination with a further hierarchical clustering were implemented to establish the brain fMRI networks. The ridge-restricted canonical correlation analysis (RCCA) was adopted to evaluate the overall correlation between the brain networks and reinforcement-related behaviours. Based on the RCCA results, we have identified the neural signatures across three brain fMRI networks for each reinforcement-related behaviour.

### Identification of Reinforcement-related Brain fMRI Networks

We defined brain networks underlying reinforcement-related behaviour by using the Monetary Incentive Delay (MID) task to measure reward processing^21^, the Stop Signal Task (SST), to assess motor inhibition^22^ and the Emotional Faces Task (EFT) to examine emotional processing^23^. In these tasks, we analysed contrasts that are most relevant to the reinforcement-related behaviour and eliciting the largest BOLD-difference, namely the ‘large win vs no win’ contrast during the reward anticipation phase in the MID task, the ‘successful stop vs successful go’ contrast in the SST, and the ‘angry face vs control’ contrast in the EFT.

We applied WVCNA ^13, 19^, which was established by combining the scale-free network assumption with a dynamic cut of the dendrogram^24^, to maximise the resolution of localised brain network features (see materials and methods for details). Using this approach, we identified in the MID a brain network consisting of 500 nodes (25130 voxels, Figure 2A); in the SST 487 nodes (24571 voxels, Figure 2B) and in the EFT 79 nodes (3923 voxels, Figure 2C). We further removed redundant information by applying an additional hierarchical clustering on these nodes with a static cut at the 90th percentile, keeping the 10% most distinctive branches (representing clusters) in each dendrogram. This two-step procedure enabled us to efficiently reduce dimensionality while still preserving localised network features from WVCNA (Table S1A-C). Using this approach, we identified 46 clusters in MID, 41 clusters in SST and 9 clusters in EFT (Table S1A-C and Figure S1A-C).

**Figure 2.**
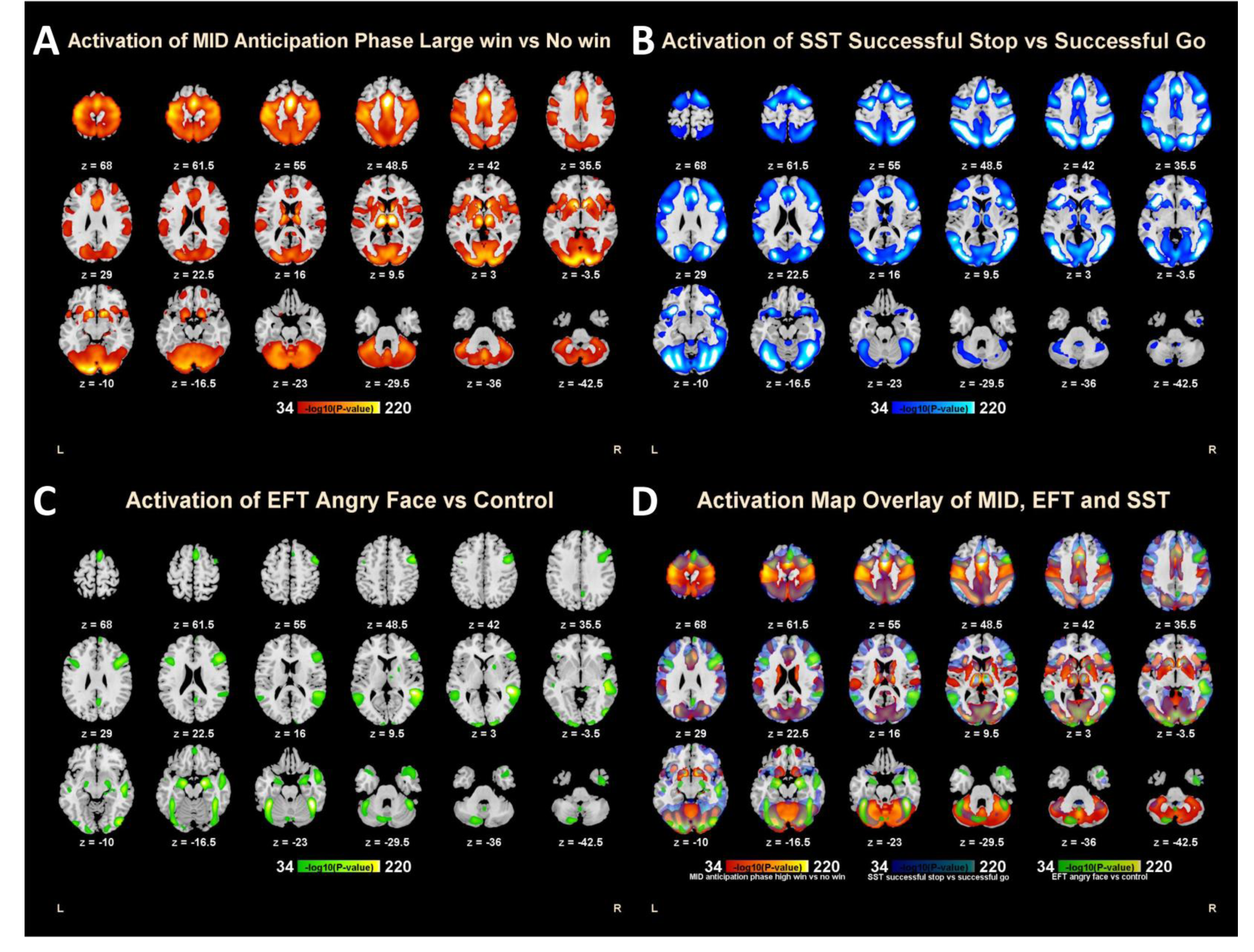
**The Activation map of MID (A), SST (B), EFT (C) and their overlay (D)**. In all figures, MID, SST and EFT were represented by red, blue and green. The activation levels were measured as the -log10 transformation of P-value and only voxels with P-value < 1.0E-34 (i.e. Effect Size D>0.3) were illustrated.

In all three networks, activated clusters were widely spread across cortical and sub- cortical regions, as well as in the cerebellum (Figure 2 and Table S1D). Brain regions activated in the three networks were often overlapping. (Figure 2D). It is notable that none of the regions of interest typically associated with reward processing or impulsiveness or emotional processing was specific to their corresponding networks. For example, VS and OFC typically linked to reward processing^9^ were activated in both MID and SST; rIFC often associated with inhibitory control^10^ was activated in both SST and EFT. STS, which is regarded as an essential component of the social brain^12^ was also activated in both SST and EFT. The dorsal amygdala, a central node of emotional processing^11^, was activated not only in EFT but also in MID. However, some activations were network-specific, for example, distinct activations during the MID in the superior post-central gyrus (i.e. the superior primary somatosensory cortex SPSC), primary auditory cortex (PAC), dorsal striatum and most of the cerebellar vermis; distinct activations in the SST were observed in the frontal operculum and the orbital part of rIFC (rIFC-Orb), inferior primary somatosensory cortex (iPSC) and the lingual part of the cerebellar vermis; and the EFT showed distinct activations in the medial orbitofrontal cortex (mOFC), dorsal posterior cingulate cortex (dPCC), temporal pole and the ventral amygdala (Figure 2D, Table S1D).

### Modulation of Reinforcement-related Brain fMRI Networks in Different Behaviours

Clinical psychopathology in adolescents is grouped into externalising and internalising disorders^25^. We were interested in examining if externalising and internalising behavioural symptoms correlate with distinct configurations of reinforcement related networks. From the Strength and Difficulties Questionnaire (SDQ) and the Development and Well-Being Assessment (DAWBA) we selected the entry-level questions, including 44 externalising items (Table 1A) covering symptoms of attentional deficit/hyperactivity disorder (ADHD; 23 items), oppositional defiance disorder (ODD; 11 items) and conduct disorder (CD; 10 items), and 21 internalising items (Table 1B) covering symptoms of depression (12 items) and anxiety (8 items) (see supplementary materials for more details). To evaluate the overall relationship of behavioural symptoms and patterns of brain activation we carried out ridge-regularised canonical correlation analysis (RCCA)^20^. This method seeks to find subsets of variables in two datasets that best correlate with each other while stabilising the result through penalisation of correlations within each dataset. We first investigated the overall correlation between externalising behaviours and 96 clusters from the three fMRI networks and found a significant canonical correlation (Hotelling’s Trace (HT)=1.86, df=(96,44), P_perm_=0.0001; see material and methods for details) (Table 2 and S2). The number of permutations to calculate p-values in this and all subsequent analyses is 10,000 unless otherwise specified. Also, presented p-values are always corrected for experimental-wise multiple comparisons wherever applicable. We then investigated the RCCA between internalising behaviours and the same 96 fMRI clusters but found no overall significance (HT=0.83, df=(96,21), P_Perm_=0.7862). We also did not find significant overall correlations with internalising behaviours when analysing each fMRI network separately (Table S3). We, therefore, constrained our subsequent analyses to externalising behaviours only.

**Table 1.**
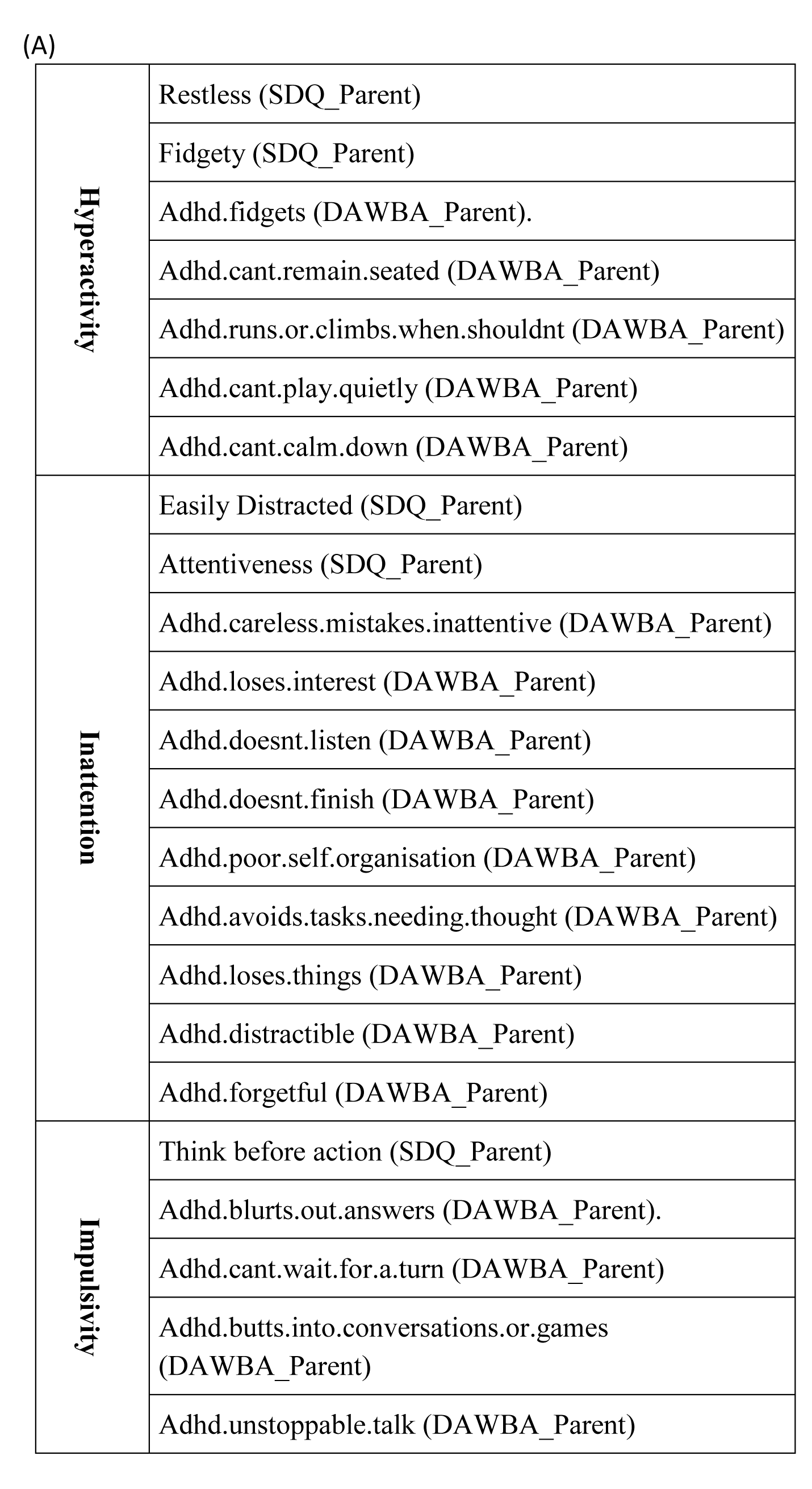
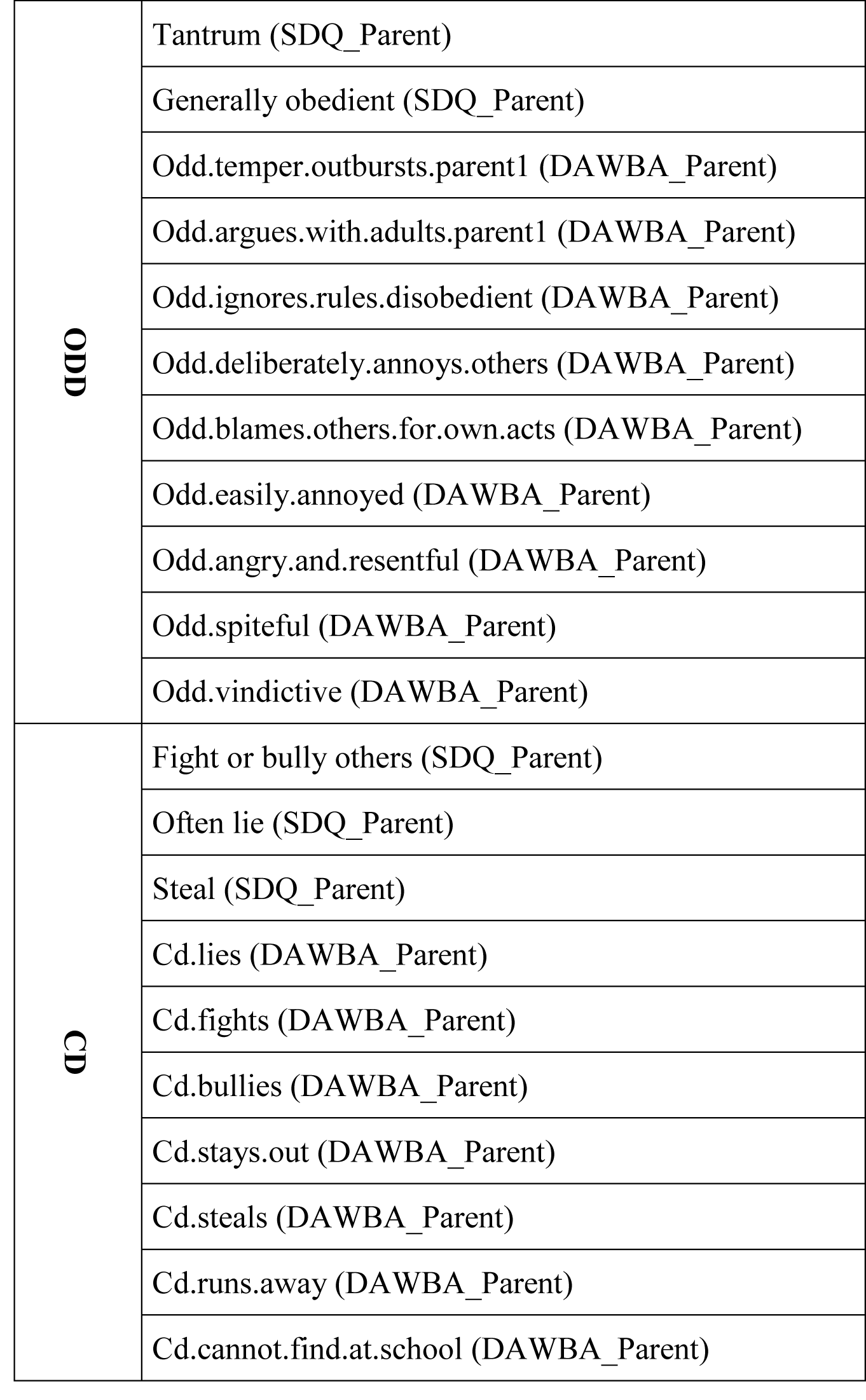
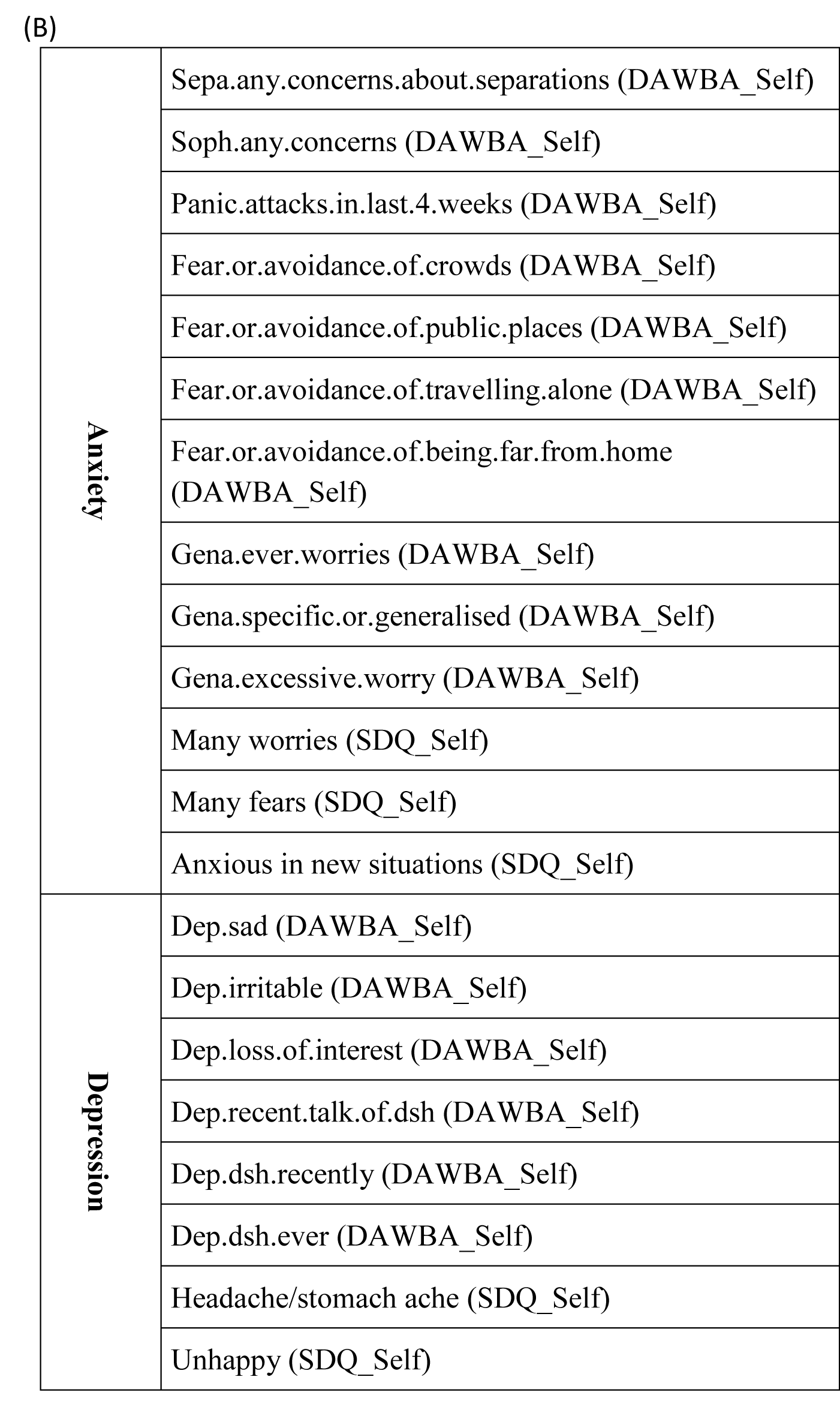
List of (A) Externalising Items from parents-rated SDQ and DAWBA, and (B) Internalising Items from child-rated SDQ and DAWBA.

|  |  |
| --- | --- |
| **Hyperactivity** | Restless (SDQ_Parent) |
|  | Fidgety (SDQ_Parent) |
|  | Adhd.fidgets (DAWBA_Parent). |
|  | Adhd.cant.remain.seated (DAWBA_Parent) |
|  | Adhd.runs.or.climbs.when.shouldnt (DAWBA_Parent) |
|  | Adhd.cant.play.quietly (DAWBA_Parent) |
|  | Adhd.cant.calm.down (DAWBA_Parent) |
| **Inattention** | Easily Distracted (SDQ_Parent) |
|  | Attentiveness (SDQ_Parent) |
|  | Adhd.careless.mistakes.inattentive (DAWBA_Parent) |
|  | Adhd.loses.interest (DAWBA_Parent) |
|  | Adhd.doesnt.listen (DAWBA_Parent) |
|  | Adhd.doesnt.finish (DAWBA_Parent) |
|  | Adhd.poor.self.organisation (DAWBA_Parent) |
|  | Adhd.avoids.tasks.needng.thought (DAWBA_Parent) |
|  | Adhd.loses.things (DAWBA_Parent) |
|  | Adhd.distractible (DAWBA_Parent) |
|  | Adhd.forgetful (DAWBA_Parent) |
| **Impulsivity** | Think before action (SDQ_Parent) |
|  | Adhd.blurts.out.answers (DAWBA_Parent). |
|  | Adhd.cant.wait.for.a.turn (DAWBA_Parent) |
|  | Adhd.butts.into.conversations.or.games (DAWBA_Parent) |
|  | Adhd.unstoppable.talk (DAWBA_Parent) |
| ODD | Tantrum (SDQ_Parent) |
|  | Generally obedient (SDQ_Parent) |
|  | Odd.temper.outbursts.parent1 (DAWBA_Parent) |
|  | Odd.argues.with.adults.parent1 (DAWBA_Parent) |
|  | Odd.ignores.rules.disobedient (DAWBA_Parent) |
|  | Odd.deliberately.annoys.others (DAWBA_Parent) |
|  | Odd.blames.others.for.own.acts (DAWBA_Parent) |
|  | Odd.easily.annoyed (DAWBA_Parent) |
|  | Odd.angry.and.resentful (DAWBA_Parent) |
|  | Odd.spiteful (DAWBA_Parent) |
|  | Odd.vindictive (DAWBA_Parent) |
| CD | Fight or bully others (SDQ_Parent) |
|  | Often lie (SDQ_Parent) |
|  | Steal (SDQ_Parent) |
|  | Cd.lies (DAWBA_Parent) |
|  | Cd.fights (DAWBA_Parent) |
|  | Cd.bullies (DAWBA_Parent) |
|  | Cd.stays.out (DAWBA_Parent) |
|  | Cd.steals (DAWBA_Parent) |
|  | Cd.runs.away (DAWBA_Parent) |
|  | Cd.cannot.find.at.school (DAWBA_Parent) |
| (B) |  |
| **Anxiety** | Sepa.any.concerns.about.separations (DAWBA_Self) |
|  | Soph.any.concerns (DAWBA_Self) |
|  | Panic.attacks.in.last.4.weeks (DAWBA_Self) |
|  | Fear.or.avoidance.of.crowds (DAWBA_Self) |
|  | Fear.or.avoidance.of.public.places (DAWBA_Self) |
|  | Fear.or.avoidance.of.travelling.alone (DAWBA_Self) |
|  | Fear.or.avoidance.of.being.far.from.home<br>(DAWBA_Self) |
|  | Gena.ever.worries (DAWBA_Self) |
|  | Gena.specific.or.generalised (DAWBA_Self) |
|  | Gena.excessive.worry (DAWBA_Self) |
|  | Many worries (SDQ_Self) |
|  | Many fears (SDQ_Self) |
|  | Anxious in new situations (SDQ_Self) |
| **Depression** | Dep.sad (DAWBA_Self) |
|  | Dep.irritable (DAWBA_Self) |
|  | Dep.loss.of.interest (DAWBA_Self) |
|  | Dep.recent.talk.of.dsh (DAWBA_Self) |
|  | Dep.dsh.recently (DAWBA_Self) |
|  | Dep.dsh.ever (DAWBA_Self) |
|  | Headache/stomach ache (SDQ_Self) |
|  | Unhappy (SDQ_Self) |

**Table 2.**
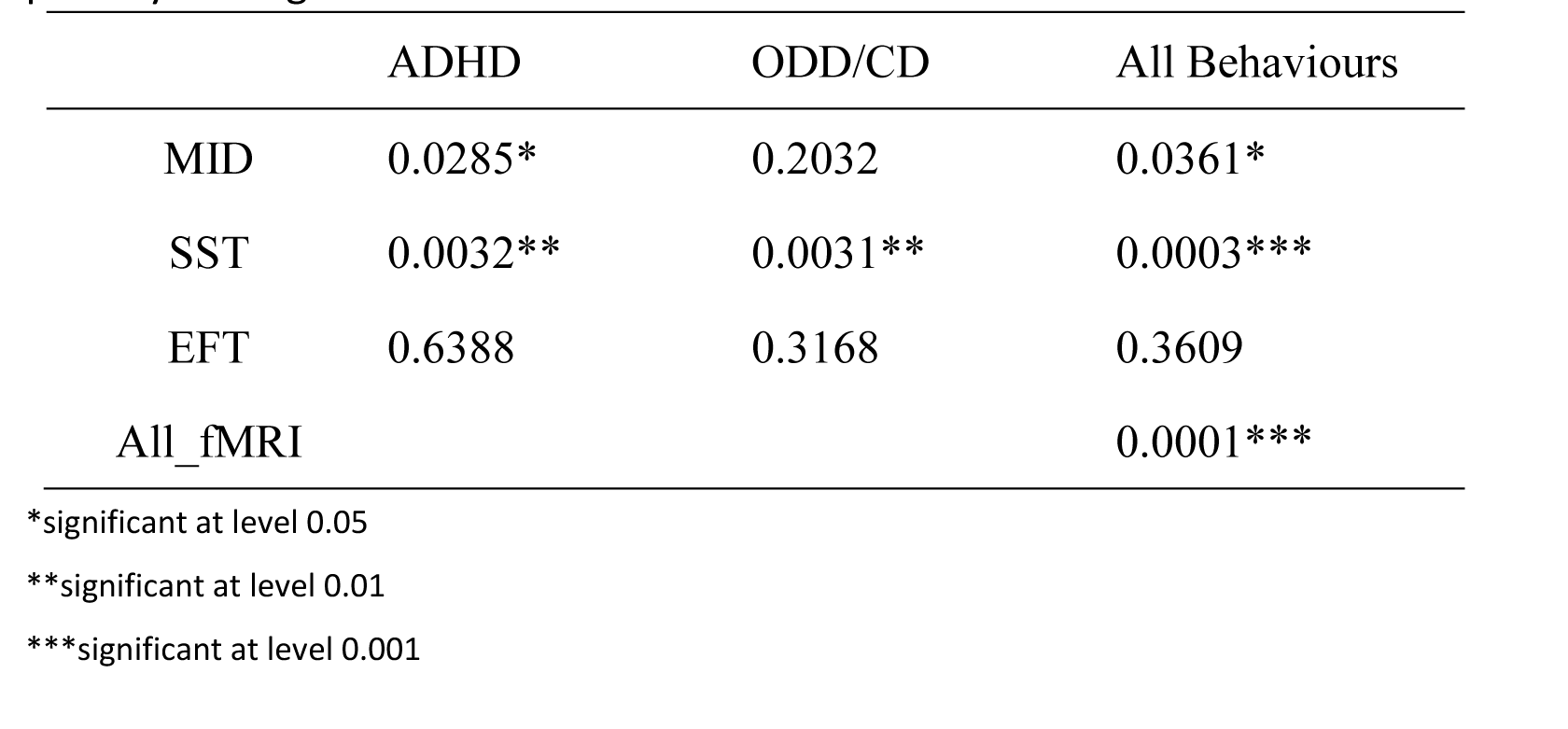
Regularised CCA P-values based on 10000 Permutation with penalty λ=0.1 for both fMRI and externalising behaviour items. Similar results have been achieved with varied penalty settings and shown in Table S2.

Next, we investigated the contribution of each brain network to different behavioural conditions. For the reward anticipation network, we found an overall significant correlation with externalising behaviours (HT=0.85, df=(46,44), P_Perm_=0.0361, Table 2 and S2). This correlation was driven by ADHD behaviours (HT=0.45, df=(46,23), P_Perm_=0.0285, Table 2 and S2), but not ODD/CD behaviours (HT=0.41, df=(46,21), P_Perm_=0.2032, Table 2 and S2), indicating that reward anticipation mainly relates to ADHD symptoms. For the motor inhibition network, we found an overall significant correlation with externalising behaviours (HT=0.83, df=(41,44), P_Perm_ =0.0003, Table 2). This correlation was driven by ADHD behaviours (HT=0.45, df=(46,23), P_Perm_=0.0032, Table 2), as well as ODD/CD behaviours (HT=0.41, df=(41,21), P_Perm_=0.0031, Table 2), indicating that motor inhibition might play a role in both ADHD and ODD/CD symptoms. For the emotional processing network, we found neither significant correlation with externalising behaviours (HT=0.19, df=(9,44), P_Perm_=0.392, Table 2 and S2), or with ADHD behaviours (HT=0.09, df=(9,23), P_Perm_=0.634, Table 2 and S2) or with ODD/CD behaviours (HT=0.10, df=(9,21), P_Perm_=0.294, Table 2 and S2) alone.

### Functional brain characterisation of behaviours across different tasks

While both reward anticipation and motor inhibition networks show significant canonical correlations with ADHD behaviours, neither first eigenvalue (the squared correlation between the first components of RCCA) was significant on its own (P_Perm_=0.0867 for reward anticipation; P_Perm_=0.1506 for motor inhibition), suggesting distinctive neural bases underlying different ADHD behaviours. Therefore, we investigated profiles across brain networks that may characterise the ADHD components hyperactivity, inattention or impulsivity (see Materials and Methods). As the factors generated by RCCA are not optimised to detect differences in the brain function underlying these behaviours, we applied a more sensitive multivariate linear model. Together, reward anticipation and motor inhibition networks were found in significant association with the summed score (i.e. the total score) of ADHD behaviours (F_(87,1418)_=1.51, P=2.23×10^−3^), as well as the total scores of ADHD components hyperactivity (F_(87,1418)_=1.58, P=6.97×10^−4^), impulsivity (F_(87,1418)_=1.37, P=0.0166) and inattention (F_(87,1418)_=1.40, P=0.0111). However, we observed differential associations of these ADHD behaviours with reward anticipation and motor inhibition networks: while the motor inhibition network was found in significant association with the total scores of all three ADHD components (F_(41,1464)_=1.67, P=5.23×10^−3^ for hyperactivity; F_(41,1464)_=1.92, P=4.77×10^−4^ for impulsivity; F_(41,1464)_=1.58, P=0.0113 for inattention), the reward anticipation network only showed a significant association with the total score of hyperactivity (F_(46,1459)_=1.427, P=0.0329), but not with those of impulsivity (F_(46,1459)_=0.885, P=0.691) and inattention (F_(46,1459)_=1.214, P=0.156).

#### fMRI signature for hyperactivity

The hyperactivity total score was significantly associated with six out of 46 brain regions in the reward anticipation network: superior parietal lobule (SPL), middle central sulcus (mid- CS), thalamus, primary auditory cortex (PAC), middle cingulate cortex (MCC) and superior frontal junction (SFJ) (Figure 3A, Table 3A and Table S4). We investigated the specificity of the observed associations and found that SPL, mid-CS and thalamus were also associated with inattention, and SPL, mid-CS and MCC were associated with ODD/CD behaviours, whereas no significant association was found with impulsivity (Table 3A and Table S4). The brain regions showed similar association strength with hyperactivity and inattention (P_Perm_=0.8337, as well as with ODD/CD behaviours (P_Perm_ =1) but significantly weaker in the case of impulsivity (P_Perm_=0.0168) (Table S5). Together, our findings suggest a shared specificity of brain activation during reward anticipation in hyperactivity, inattention and ODD/CD behaviours, but not in impulsivity (Figure 3E).

**Figure 3.**
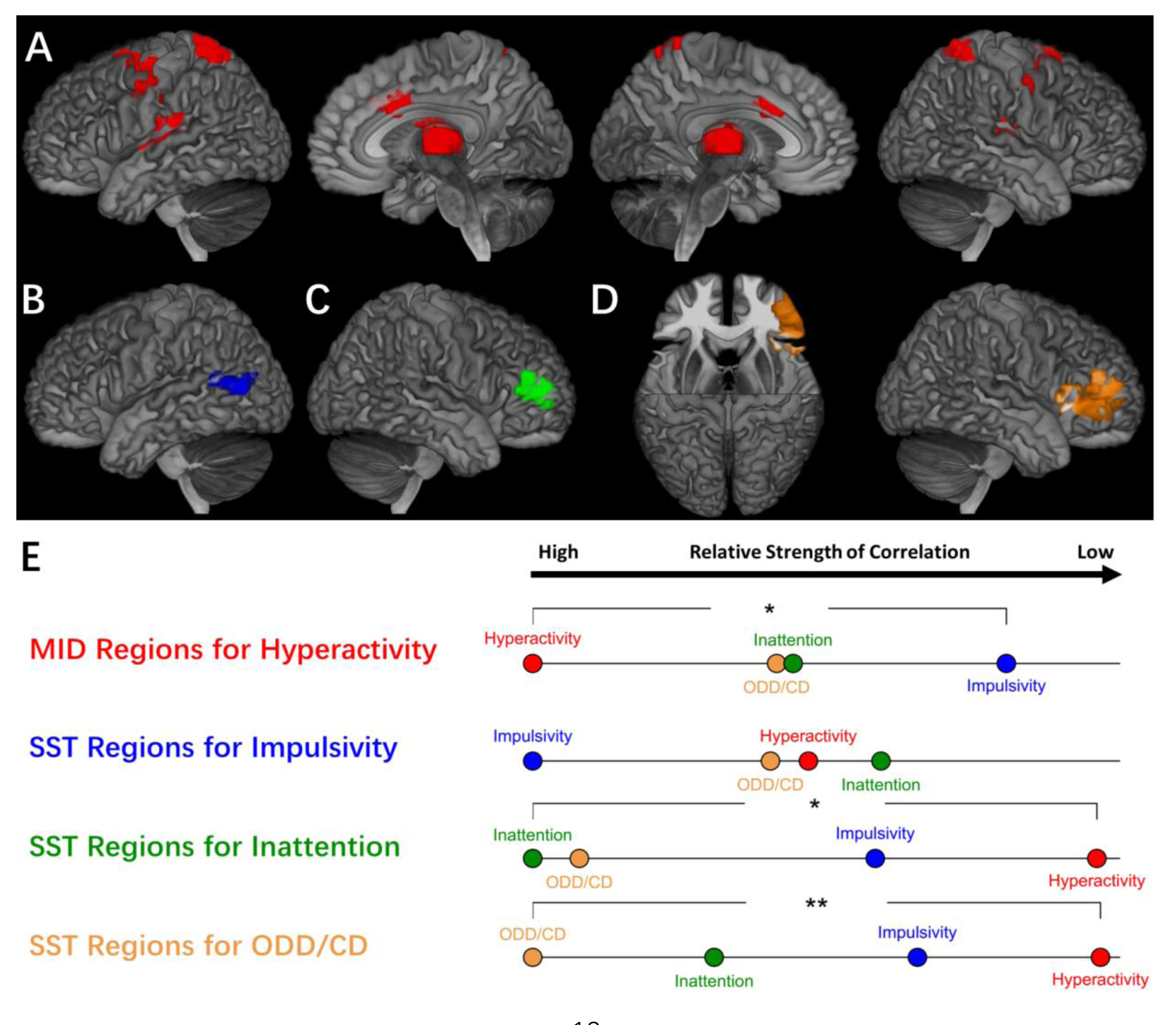
A. Reward anticipation network underlying Hyperactivity. (Red: Thalamus, Superior Parietal Lobule, middle Central Sulcus, Primary Auditory Cortex, Middle Cingulate Cortex and Superior Frontal Junction); **B. Motor inhibition network underlying Impulsivity** (Blue: left middle Temporal-Parietal Junction); **C. Motor inhibition network underlying Inattention** (Green: right anterior Inferior Frontal Sulcus); **D. Motor inhibition network underlying ODD/CD behaviours** (Yellow: right Inferior Frontal Gyrus + anterior Insula and right anterior Inferior Frontal Sulcus); **E. Neural signatures of ADHD and ODD/CD behaviours.** For each neural network identified in A-D, its correlations with the corresponding primary behaviour and the rest ADHD or ODD/CD behaviours were compared that the corresponding relative strength of correlations were plotted (Red: Hyperactivity; Blue: Impulsivity; Green: Inattention; Yellow: ODD/CD behaviours). Pairwise significant difference in correlations were highlighted with ‘*’ (significant at 0.05 level) or ‘**’ (significant at 0.01 level).

**Table 3.** Prominent clusters of brain networks for (A) hyperactivity, (B) impulsivity, (C) inattention and (D) ODD/CD behaviours. For each behaviour component, the prominent clusters in each brain network were identified if their correlations with the corresponding behaviour component (i.e. column ‘Primary Behaviour’) were significant after correction for multiple comparison (10000-permutation) and were highlighted in bold as well as with sign † (column Pcorrected). For all prominent clusters identified in the first step, we further explored their correlations with the remaining behaviour components (i.e. column ‘Exploratory Analyses), where significant correlations after correction for multiple testing were highlighted in bold as well as with either sign * or ** representing the significance at level 0.05 and 0.01, respectively. ‡ These one-tailed P-values were based on 10000-permutation after correction for the number of corresponding exploratory tests, i.e. 18 for (A), 3 for (B), 3 for (C) and 6 for (D). See Table S4, S6-7 for the complete results.

**A**
| MID Regions | Primary Behaviour |  | Exploratory Analyses |  |  |  |  |  |
| --- | --- | --- | --- | --- | --- | --- | --- | --- |
|  | Hyperactivity |  | Impulsivity |  | Inattention |  | ODD/CD |  |
| | R | $P_{\text{corrected}}$ | R | $P_{\text{one-tailed}}^{\ddagger}$ | R | $P_{\text{one-tailed}}^{\ddagger}$ | R | $P_{\text{one-tailed}}^{\ddagger}$ |
| Thalamus | **-0.091** | **0.0113†** | -0.032 | 0.5447 | **-0.074** | **0.0241\*** | -0.065 | 0.0626 |
| SFJ | **-0.084** | **0.0285†** | -0.014 | 0.8547 | -0.066 | 0.0566 | -0.052 | 0.1796 |
| PAC | **-0.085** | **0.0250†** | -0.025 | 0.6731 | -0.051 | 0.2025 | -0.063 | 0.0711 |
| SPL | **-0.094** | **0.0072†** | -0.045 | 0.2863 | **-0.077** | **0.0174\*** | **-0.068** | **0.0483\*** |
| mid-CS | **-0.091** | **0.0110†** | -0.040 | 0.3802 | **-0.075** | **0.0240\*** | **-0.074** | **0.0243\*** |
| MCC | **-0.084** | **0.0270†** | -0.017 | 0.8068 | -0.046 | 0.2696 | **-0.078** | **0.0164\*** |

**B**
| SST Regions | Primary Behaviour |  | Exploratory Analyses |  |  |  |  |  |
| --- | --- | --- | --- | --- | --- | --- | --- | --- |
|  | Impulsivity |  | Hyperactivity |  | Inattention |  | ODD/CD |  |
| | R | $P_{\text{corrected}}$ | R | $P_{\text{one-tailed}}^{\ddagger}$ | R | $P_{\text{one-tailed}}^{\ddagger}$ | R | $P_{\text{one-tailed}}^{\ddagger}$ |
| Left TPJ | **-0.092** | **0.0093†** | **-0.067** | **0.0142\*** | **-0.058** | **0.0347\*** | **-0.071** | **0.00895\*\*** |

**C**
| SST Regions | Primary Behaviour |  | Exploratory Analyses |  |  |  |  |  |
| --- | --- | --- | --- | --- | --- | --- | --- | --- |
|  | Inattention |  | Hyperactivity |  | Impulsivity |  | ODD/CD |  |
| | R | $P_{\text{corrected}}$ | R | $P_{\text{one-tailed}}^{\ddagger}$ | R | $P_{\text{one-tailed}}^{\ddagger}$ | R | $P_{\text{one-tailed}}^{\ddagger}$ |
| Right aIFS | **-0.087** | **0.0191†** | -0.017 | 0.758 | **-0.056** | **0.0434\*** | **-0.084** | **0.00170\*\*** |

**D**
| SST Regions | Primary Behaviour |  | Exploratory Analyses |  |  |  |  |  |
| --- | --- | --- | --- | --- | --- | --- | --- | --- |
|  | ODD/CD |  | Hyperactivity |  | Impulsivity |  | Inattention |  |
| | R | $P_{\text{corrected}}$ | R | $P_{\text{one-tailed}}^{\ddagger}$ | R | $P_{\text{one-tailed}}^{\ddagger}$ | R | $P_{\text{one-tailed}}^{\ddagger}$ |
| **Right IFC +**<br>**aInsula** | **-0.090** | **0.0111†** | -0.014 | 1 | -0.045 | 0.238 | -0.053 | 0.115 |
| **Right aIFS** | **-0.084** | **0.0270†** | -0.017 | 1 | -0.056 | 0.0867 | **-0.087** | **0.00207\*\*** |

In the motor inhibition network, however, despite the overall significant association, none of the six brain regions was significantly associated with hyperactivity (Table S6A), suggesting the observed overall association was based on multiple fMRI regions of the motor inhibition network, each with a minor contribution.

#### fMRI signature for impulsivity

The left temporoparietal junction (TPJ) of the motor inhibition network was associated with impulsivity (R=-0.092, t=-3.563, P_Perm_=0.0097) (Figure 3B, Table 3B and Table S6B), and additionally in exploratory association with hyperactivity (R=-0.067, P_one-tailed_=0.0142), inattention (R=−0.058, P_one-tailed_=0.0347) and ODD/CD behaviours (R=-0.071, P_one-tailed_=8.95×10^−3^) (Table 3B and Table S6B) with similar association strength (P_Perm_ =0.8229, P_Perm_=0.4559 and P_Perm_ =1). Together, this suggests a shared specificity across ADHD and ODD/CD behaviours during motor inhibition (Figure 3E).

#### fMRI signature for inattention

In the motor inhibition network, we found significant association of the right anterior inferior frontal sulcus (aIFS) with inattention (R=-0.087, t=-3.368, P_Perm_=0.0191), as well as with impulsivity (R=-0.056, P_one-tailed_=0.0434) and ODD/CD behaviours (R=-0.084, P_one-tailed_=1.70×10^−3^), but not with hyperactivity (R=-0.017, P_one-tailed_=0.758) (Figure 3C, Table 3C and Table S6C). The strength of association of aIFS with inattention is comparable to impulsivity (P_Perm_=0.5622) and ODD/CD behaviours (P_Perm_=1), but significantly stronger than that with hyperactivity (P_Perm_=0.0168), suggesting distinct specificities of hyperactivity and inattention during motor inhibition, and shared specificity of inattention with impulsivity and ODD/CD behaviours (Figure 3E).

#### fMRI signatures for ODD/CD behaviours

ODD/CD behaviours were found only in a significant canonical correlation with the motor inhibition network. Right aIFS (R=-0.084, t=-3.26, P_Perm_=0.0270) and right IFC/anterior insula (R=-0.090, t=-3.51, P_Perm_=0.0111) were associated with the summed score of ODD/CD behaviours (Figure 3D, Table 3D and Table S7). Both associations were mainly driven by the ODD behaviours (t=-3.64, P_one-tailed_=1.43×10^−4^ for the right aIFS; t=−3.77, P_one-tailed_=8.34×10^−5^ for the right IFC/anterior insula), and less by the CD behaviours (t=−1.775, P_one-tailed_=0.0380 for the right aIFS; t=−2.12, P_one-tailed_=0.0170 for the right IFC/anterior insula) (Table S7). Together ODD/CD prominent regions showed similar association strength with ODD/CD behaviours and Inattention (P_Perm_=1), as well as with impulsivity (P_Perm_=0.2742), but significantly lower with hyperactivity (P_Perm_=0.0072) (Figure 3E and Table S8).

In conclusion of the above results, ADHD and ODD/CD may share several distinctive neural bases during reward anticipation and motor inhibition.

## Discussion

Here we characterise clinically relevant behaviours in adolescents by describing brain activation during reinforcement-related cognitive processes. These behaviours include externalising symptoms of hyperactivity, impulsiveness and inattention, oppositional defiance and conduct, and internalising symptoms of anxiety and depression. We interrogate the neural basis of each of these behaviours by measuring brain activity during reinforcement-related neuroimaging tasks of reward processing, motor inhibition and emotional processing.

### Shared and distinct activation patterns across reinforcement-related tasks

We find that activation of similar brain regions is often associated with different tasks (and behaviours). While well-known representative brain areas (e.g. VS and OFC for reward anticipation^9^, right-IFC for inhibitory control^10^, and amygdala and STS for emotional processing ^11, 12^) were activated as expected, these activations were not restricted to one task alone. This might represent the involvement of shared cognitive components in different behaviours that might be less specific to individual tasks. For example, the VS activation during motor inhibition was due to the anticipation of a random event^26^, thus sharing the anticipatory component with the reward anticipation network that also activates the same region. In some instances, it may also be caused by brain activation that reflects task presentation (for example, motor cortex activation in the ‘active’ MID and SSRT, but not in the passive viewing EFT). Our observation is consistent with the notion of a basic neural function that underlies a complex profile of different behaviours^27^.

However, the overlap of brain activation across neuroimaging tasks might also indicate the presence of different functional or structural domains within a given brain region that relate differentially to each task^28^. This latter hypothesis is supported by the observation of low correlations of the same brain regions across tasks. In contrast, we found high correlations between different brain regions within each task, suggesting network constellations that are specific to each individual neuroimaging task. This specificity was further suggested by the observation that the variance of hyperactivity explained by reward anticipation and motor inhibition networks are additive, and thus not overlapping. Specificity of cognitive neural networks might thus be defined as much by their internal collaborative structure as by the individual brain regions involved^29^.

### Activation of sensory cortical brain regions is not specific to the quality of the stimulus

We found regions solely activated in the MID task, which hitherto had not been implicated in anticipation of a visually presented reward. They included, for instance, the primary auditory cortex (PAC) that we observed to be activated in the absence of any auditory stimulus. As the PAC has been found to predict reward value^17^ and is associated with anticipatory motor response^30^ upon auditory stimulation, our findings point towards the possibility of the PAC underlying these cognitive processes in a way that is not dependent on the quality of the sensory stimulus. In addition, wide areas within the somatosensory cortex were solely activated in the MID task, further suggesting the recruitment of sensory cortices (including the visual cortices) during reward anticipation irrespective of the quality of the signal input^31^.

### Stratification of behavioural components by neural signatures

We found a strong overall correlation of neural networks with externalising behaviours (ADHD and ODD/CD), particularly in reward anticipation and motor inhibition, but no significant correlation with internalising behaviours. This discrepancy might be due to the young age of the participants, as at 14 years the prevalence of internalising disorders (e.g. depression) is still accumulating. Furthermore, internalising disorders have a lower prevalence than externalising disorders^32^, reducing the power of our sample to detect their correlation with neural networks.

While ADHD behaviours were related to both reward anticipation and motor inhibition networks, we found specific neural signatures that distinguished each of the individual behaviours. While brain activity in the reward anticipation network was correlated with both hyperactivity and inattention, their activation patterns were not significantly different (Figure 3 and Table S4&S5). However, in the motor inhibition network, the correlation with inattention was significantly higher than that with hyperactivity (Figure 3 and Table S6C), consistent with a greater effort to maintain sustained attention during the task. This interpretation is supported by the strong correlation during successful motor inhibition of inattention with right inferior frontal cortical activity, a brain region previously implicated in attentional detection, monitoring and motor inhibition^10^.

In contrast, in impulsivity we found no correlation with the reward anticipation network. In the motor inhibition network, its strongest correlation was with activation of the left temporal-parietal junction (TPJ), but there were no significant differences to the activation patterns of both hyperactivity and inattention (Figure 3 and Table S6B). This observation is in line with the previous finding of reduced bilateral TPJ activity in ADHD patients^33^.

We, thus, identify neural signatures that distinguish hyperactivity, inattention and impulsivity on the basis of brain activation patterns during reward anticipation and motor inhibition. These signatures enable a more refined characterisation of ADHD behaviour than the currently used distinction between motivational vs motor inhibitory processes^34^.

### Biological characterisation of comorbidity by neural signatures

ODD/CD behaviours were only related to the motor inhibition network, but not reward anticipation, which is in line with previous findings ^35, 36^. Activation patterns for ODD and CD behaviours in the motor inhibition network were similar, suggesting a shared neural basis (Figure 3 and Table S7)^37^. Surprisingly, we were not able to distinguish activation patterns in the motor inhibition network in conduct and inattention symptoms. While this may indicate in part a shared neural basis, the phenotypic differences between these behaviours also suggest the presence of a distinguishing cognitive domain, which we have not captured in our tasks. Nevertheless, the shared neural signatures between ODD/CD and ADHD symptoms indicate a shared neural basis underlying the high comorbidity between ODD/CD and ADHD^38, 39^, supporting the idea of unifying ADHD and ODD/CD into a single spectrum disorder^40^.

Our approach provides a unified framework to investigate brain activity in reinforcement-related behaviour enabling the characterisation of shared and distinct functional brain activation patterns that underlie different externalising symptoms. It also results in the identification of neural signatures that may help to stratify these symptoms, while accounting for clinically observed co-morbidity.

## Materials and Methods

### fMRI Data Acquisition and Analysis

Structural and functional MRI data were acquired at eight IMAGEN assessment sites with 3T MRI scanners of different manufacturers (Siemens, Philips, General Electric, Bruker). The scanning variables were specifically chosen to be compatible with all scanners. The same scanning protocol was used in all sites. In brief, high-resolution T1-weighted 3D structural images were acquired for anatomical localization and co-registration with the functional time-series. Blood-oxygen-level-dependent (BOLD) functional images were acquired with gradient-echo, echo-planar imaging (EPI) sequence. For the MID task, 300 volumes were acquired for each participant, and each volume consisted of 40 slices aligned to the anterior commission/posterior commission line (2.4 mm slice thickness, 1 mm gap). The echo-time (TE) was optimized (TE=30 ms, repetition time (TR)=2,200 ms) to provide reliable imaging of subcortical areas.

Functional MRI data were analysed with SPM8 (Statistical Parametric Mapping, http://www.fil.ion.ucl.ac.uk/spm). Spatial preprocessing included: slice time correction to adjust for time differences due to multi-slice imaging acquisition, realignment to the first volume in line, non-linearly warping to the MNI space (based on a custom EPI template (53×63×46 voxels) created out of an average of the mean images of 400 adolescents), resampling at a resolution of 3×3×3mm^3^ and smoothing with an isotropic Gaussian kernel of 5 mm full-width at half-maximum.

At the first level of analysis, changes in the BOLD response for each subject were assessed by linear combinations at the individual subject level, for each experimental condition (e.g. reward anticipation high gain of Monetary Incentive Delay (MID) task), each trial was convolved with the hemodynamic response function to form regressors that account for potential noise variance, e.g. head movement, associated with the processing of reward anticipation. Estimated movement parameters were added to the design matrix in the form of 18 additional columns (three translations, three rotations, three quadratic and three cubic translations, and every three translations with a shift of ±1 TR).

For the MID anticipation phase we contrasted brain activation during ‘anticipation of high win [here signaled by a circle] vs anticipation of no-win [here signaled by a triangle]’; For the emotional faces task (EFT) we contrasted brain activation during ‘viewing Angry Face vs viewing Control [circles]’; For the stop signal task (SST) we contrasted brain activation during ‘successful stop vs successful go’. The single-subject contrast images were then taken to the population-based weighted co-activation network analysis.

### The Monetary Incentive Delay Task for fMRI

Participants performed a modified version of the Monetary Incentive Delay (MID) task to examine neural responses to reward anticipation and reward outcome^21^. The task consisted of 66 10-second trials. In each trial, participants were presented with one of three cue shapes (cue, 250 ms) denoting whether a target (white square) would subsequently appear on the left or right side of the screen and whether 0, 2 or 10 points could be won in that trial. After a variable delay (4,000-4,500 ms) of fixation on a white crosshair, participants were instructed to respond with left/right button-press as soon as the target appeared. Feedback on whether and how many points were won during the trial was presented for 1,450 ms after the response (Figure S2). Using a tracking algorithm, task difficulty (i.e. target duration varied between 100 and 300 ms) was individually adjusted such that each participant successfully responded on ∼66% of trials. Participants had first completed a practice session outside the scanner (∼5 minutes), during which they were instructed that for each 5 points won they would receive one food snack in the form of small chocolate candies.

Based on prior research suggesting reliable associations between ADHD-symptoms and fMRI BOLD responses measured during reward anticipation, the current study used the contrast ‘anticipation of high-win vs anticipation of no-win’. Only successfully ‘hit’ trials were included here.

### The Emotional Reactivity fMRI Paradigm (Emotional Faces Task)

This task was adapted from^23^. Participants watched 18-second blocks of either a face movie (depicting anger or neutrality) or a control stimulus. Each face movie showed black and white video clips (200-500ms) of male or female faces. Five blocks each of angry and neutral expressions were interleaved with nine blocks of the control stimulus. Each block contained eight trials of 6 face identities (3 female). The same identities were used for the angry and neutral blocks. The control stimuli were black and white concentric circles expanding and contracting at various speeds that roughly matched the contrast and motion characteristics of the face clips (Figure S3).

The neutral blocks contained emotional expressions that were not attributable to any particular emotion (e.g. nose twitching); however previous research has suggested that neutral stimuli are not always interpreted as such. Functional imaging studies have found significant activation of the amygdala in response to the presentation of neutral faces in healthy adult males^41^, social anxiety patients and matched control participants^42^, adolescents with conduct disorder problems^43^ and young men with violent behaviour problems^44^. This suggests that neutral faces may be interpreted as emotionally ambiguous. This study focused specifically on the effects of viewing angry faces (vs control) to eliminate this ambiguity so that any significant relationships between behaviour and brain could be interpreted as the consequence of viewing negative social stimuli (anger).

### The Stop Signal Task for fMRI

Participants performed an event-related stop signal task (SST) task designed to study neural responses to successful and unsuccessful inhibitory control^22^. The task was composed of Go trials and Stop trials. During Go trials (83%; 480 trials) participants were presented with arrows pointing either to the left or to the right. During these trials, subjects were instructed to make a button response with their left or right index finger corresponding to the direction of the arrow. In the unpredictable Stop trials (17%; 80 trials), the arrows pointing left or right were followed (on average 300 ms later) by arrows pointing upwards; participants were instructed to inhibit their motor responses during these trials (Figure S4). A tracking algorithm changes the time interval between Go signal and Stop signal onsets according to each subject’s performance on previous trials (average percentage of inhibition over previous Stop trials, recalculated after each Stop trial), resulting in 50% successful and 50% unsuccessful inhibition trials. The inter-trial interval was 1,800 ms. The tracking algorithm of the task ensured that subjects were successful on 50% of Stop trials and worked at the edge of their own inhibitory capacity.

### Population-based Weighted Voxel Co-activation Network Analysis

The weighted voxel co-activation network analysis (WVCNA)^13, 19^ was applied to parcellate those highly co-activated voxels in all three fMRI contrasts, e.g. large win vs no win contrast anticipation phase of MID task, angry face vs control contrast of face task and successful stop vs successful go contrast of SST. Such a parcellation procedure could effectively reduce the dimensionality without losing too much information. The procedure is summarised as below:

**Pre-processing.** For all three tasks, the initial pre-processing steps involved removing null voxels (including the removal of out-brain voxels based on Automated Anatomical Labelling (AAL) template) and potential participant outliers from contrast data based on low inter-sample correlations. The activation maps of pre-processed data were then generated and only those positive activations with at least a median effect size, i.e. Cohen’s D>0.3, will be included in the following analyses.

**Parameter Selection.** To minimize the arbitrary choice of parameters, we took the default and suggested settings of R package ‘WGCNA’^45^, except for the soft-thresholds of adjacency matrices, which were determined as 7 for the MID, 8 for the EFT and 7 for the SST based on the fitness of scale free topology criteria (Figure S5 A-C). The above adjacency matrices will then be used to generate the topology overlapping matrices (TOMs), which capture both the direct and indirect connections among voxels. The hierarchical clustering will then be applied on the distance matrices, as 1-TOMs, and together with the dynamic cut tree function, the fMRI modules will be generated as functional ROIs. The first principle component of each module will be included in the following analysis to represent the brain activation (or BOLD response). No merge of modules will be conducted after the hierarchical clustering to avoid using an arbitrary threshold.

### Regularised Canonical Correlation Analysis (RCCA)

CCA has been widely used to investigate the overall correlation between two sets of variables^46^. However, in our case, due to high intra-correlations in both brain fMRI networks and behavioural items, multicollinearity is a potential risk factor that could jeopardise the validity of following statistical inference. Therefore, we will adopt the ridge regularised canonical correlation proposed by^20^, where two ridge regulation parameters, λx and λy, will be added to the diagonals of corresponding covariance matrices to avoid the singularity.

As our purposes are not to maximise the power of prediction, instead of estimating the optimal regulation parameters^47^, we will fix the regulation parameters across all analyses. Although varied regulation parameters have been experimented, i.e. 0.1, 0.15, 0.2 and 0.3 for both λ, the significance of major results are consistent throughout all settings (Table S2), and therefore we will simply report the P-values as well as relevant statistics based on regulation parameter 0.1. It is also noteworthy that the optimisation of regulation parameter will almost surely invalid any attempt of calculating internalised P-values through permutation test unless the optimisation procedure is also permuted, which is very difficult, if not impossible, due to the extremely high computational demanding of optimisation at each iteration. It should also be noted that current optimisation procedures of CCA related approach focus on maximising the prediction power for the first component and therefore is not a ‘real’ optimum for our purpose of evaluating the overall correlation described below.

To evaluate the significance of RCCA, the P-values or significance level will be determined through permutation tests, where individual IDs of behaviour items will be randomly shuffled at each iteration to generate the null distribution of statistics of interest. Particularly, we use the sum of eigenvalues (i.e. the squared canonical correlations) as the test statistic, which is also known as the Hotelling’s trace^48^.

### Strength Difficulty Questionnaire (SDQ) and DAWBA

The Strength and Difficulties Questionnaire (SDQ)^49^ is a brief 25-item behavioural screening tool probing hyperactivity, emotional symptoms, conduct problems, peer problems and prosocial behaviour for 3-16 years old. In the current study, we chose parent-rated hyperactivity (5 items) and conduct problem (5 items) to present externalising problems (Table 1A), and child-rated emotional problem (5 items) to represent internalising problems (Table 1B). Such a choice is based on findings that externalising problems scores from parents is more reliable than those from children themselves, and vice versa^50^.

In DAWBA^51^, similar to SDQ, parents-rated ADHD and ODD/CD items (Table 1A), and child-rated special phobia, social phobia, general anxiety, fear and depressions items (Table 1B) are included in the analyses.

### Comparison of Related Associations/Correlations though Permutation

To compare two correlations, a fisher’s transformation is normally applied to firstly normalise the distributions of correlations. The transformed correlations, now follow the normal distribution, could then be directly compared, and the corresponding difference should also follow a normal distribution^52^. However, estimation for the variance of such a difference should properly count in the relationship of variables involved in calculating the correlations. For example, in the present paper, we are interested in the difference between two correlations that share one variable in common, i.e. in the form of cor(A,B) vs cor(A,C). While the analytical solution of the variance estimation for the above case has been extensively investigated in the past ^53-55^, we will additionally implement the permutation process to empirically investigate the variance, which not only is known to be robust even if the normality assumption has been violated, but also enable us to investigate multiple comparisons altogether, where the variance of summed absolute differences under the null hypothesis could be directly estimated through the permutation process.

In the present paper, we directly calculate the P-value (which is determined by the underlying variance) of the observed summed absolute difference through a permutation process as the chance of randomly observing (i.e. at each permutation iteration) a summed absolute difference larger than the original observation. For the comparison purpose, we also include the results from Steiger’s test^54^ in the supplementary tables, which are high similar to results using the permutation test.

## Acknowledgments

This work received support from the following sources: the European Union-funded FP6 Integrated Project IMAGEN (Reinforcement-related behaviour in normal brain function and psychopathology) (LSHM-CT- 2007-037286), the Horizon 2020 funded ERC Advanced Grant ‘STRATIFY’ (Brain network based stratification of reinforcement-related disorders) (695313), ERANID (Understanding the Interplay between Cultural, Biological and Subjective Factors in Drug Use Pathways) (PR-ST-0416-10004), BRIDGET (JPND: BRain Imaging, cognition Dementia and next generation GEnomics) (MR/N027558/1), the FP7 projects IMAGEMEND(602450; IMAging GEnetics for MENtal Disorders) and MATRICS (603016), the Innovative Medicine Initiative Project EU-AIMS (115300-2), the Medical Research Council Grant ’c-VEDA’ (Consortium on Vulnerability to Externalizing Disorders and Addictions) (MR/N000390/1), the Swedish Research Council FORMAS, the Medical Research Council, the National Institute for Health Research (NIHR) Biomedical Research Centre at South London and Maudsley NHS Foundation Trust and King’s College London, the Bundesministeriumfür Bildung und Forschung (BMBF grants 01GS08152; 01EV0711; eMED SysAlc01ZX1311A; Forschungsnetz AERIAL 01EE1406A, 01EE1406B), the Deutsche Forschungsgemeinschaft (DFG grants SM 80/7-2, SFB 940/2), the Medical Research Foundation and Medical research council (grant MR/R00465X/1), the Human Brain Project (HBP SGA 2). Further support was provided by grants from: ANR (project AF12-NEUR0008-01 - WM2NA, and ANR-12-SAMA-0004), the Fondation de France, the Fondation pour la Recherche Médicale, the Mission Interministérielle de Lutte-contre-les-Drogues-et-les-Conduites-Addictives (MILDECA), the Assistance-Publique-Hôpitaux-de-Paris and INSERM (interface grant), Paris Sud University IDEX 2012; the National Institutes of Health, Science Foundation Ireland (16/ERCD/3797), U.S.A. (Axon, Testosterone and Mental Health during Adolescence; RO1 MH085772-01A1), and by NIH Consortium grant U54 EB020403, supported by a cross-NIH alliance that funds Big Data to Knowledge Centres of Excellence; the 111 Project (B18015); the NSFC (81801773, 81873909); The Key Project of Shanghai Science and Technology Innovation Plan (16JC1420402); the Shanghai Municipal Science and Technology Major Project (No.2018SHZDZX01); ZHANGJIANG LAB; and the Shanghai Pujiang Project (18PJ1400900).

## Author Contributions

<u>Design of Study</u>: G.S. and T.J.

<u>Manuscript Writing and Editing</u>: T.J. and G.S. wrote the manuscript; A.I., E.B.Q, N.T., Q.L. and

B.F. edited the first draft; all authors critically reviewed the manuscript

<u>Study Principal Investigators</u>: T.B., G.B., A.L.W.B., U.B., C.B., S.D., J.F., H.F., A.G., H.G., P.G.,

A.H., B.I., J-L.M., M-L.P.M., F.N., T.P., L.P., J.H.F., M.N.S., H.W., R.W., G.S.

<u>Data acquisition</u>: E.B.Q., T.B., G.B., A.L.W.B., U.B., C.B., H.F., A.G., H.G., P.G., A.H., B.I., J-L.M.,

M-L.P.M., F.N., D.P.O., T.P., L.P., J.H.F., M.N.S., H.W., R.W., G.S.

<u>Data analysis</u>: T.J. and A.I.

